# RNA matchmaking remodels lncRNA structure and promotes PRC2 activity

**DOI:** 10.1101/2020.04.13.040071

**Authors:** Maggie M. Balas, Erik W. Hartwick, Chloe Barrington, Justin T. Roberts, Stephen K. Wu, Ryan Bettcher, April M. Griffin, Jeffrey S. Kieft, Aaron M. Johnson

## Abstract

Human Polycomb Repressive Complex 2 (PRC2) catalysis of histone H3 lysine 27 methylation at certain loci depends on long noncoding RNAs (lncRNAs). Yet, in apparent contradiction, RNA is a potent catalytic inhibitor of PRC2. Here we show that intermolecular RNA-RNA interactions between the lncRNA HOTAIR and its target genes can relieve RNA inhibition of PRC2. RNA matchmaking is promoted by heterogenous nuclear ribonucleoprotein (hnRNP) B1, which uses multiple protein domains to bind regions of HOTAIR via multi-valent protein-RNA interactions. Chemical probing demonstrates that RNA matchmaking changes HOTAIR RNA structure. Genome-wide HOTAIR/PRC2 activity occurs at genes whose transcripts can make favorable RNA-RNA interactions with HOTAIR. We demonstrate that RNA-RNA matches of HOTAIR with target gene RNAs can relieve the inhibitory effect of a single lncRNA for PRC2 activity. Our work highlights an intrinsic switch that allows PRC2 activity in specific RNA contexts, which could explain how many lncRNAs work with PRC2.

## INTRODUCTION

Chromatin regulation can depend on long noncoding RNA (lncRNA) transcripts(1), though in many cases the exact mechanism of action for the RNA is unclear. The most well-established example of lncRNA regulation of chromatin is via the Xist RNA that is required for silencing of one copy of the X chromosome in female mammals(2). Another more recent example of a lncRNA associated with chromatin-based silencing is the HOTAIR transcript(3). Both of these lncRNAs are associated with the activity of the histone methyltransferase complex Polycomb Repressive Complex 2 (PRC2), which is involved in facultative heterochromatin formation(4). The chromatin of the inactive X chromosome and of HOTAIR-repressed genes is marked in a lncRNA-dependent manner by histone H3 lysine 27 trimethylation, the product of PRC2. PRC2 binds to single stranded RNA (ssRNA) with low specificity(5), perhaps sampling nascent transcripts on chromatin(6) due to relatively fast on and off rates for RNA binding(7). When RNA binds to PRC2, the methyltransferase activity for nucleosomes is inhibited(8,9), suggesting that some specific additional context is required for PRC2 activity when RNAs such as Xist or HOTAIR are associated at chromatin that is methylated by PRC2.

Specificity of PRC2 activity is multi-faceted and differs depending on the organism and whether initiating or maintaining heterochromatin. In *Drosophila*, PRC2 is largely dependent on a small set of specific DNA binding proteins to recruit the complex for initiation of heterochromatin (*de novo* methylation)(4). Maintenance of H3K27me3 is facilitated by the ability of the EED subunit of PRC2 to recognize H3K27me3, recruiting and stimulating PRC2 at previously-established heterochromatin regions(4). This mechanism is thought to play a role in inheritance of methylation after cell division, though DNA binding proteins are still essential in this maintenance mechanism as well(4). In humans, PRC2 recruitment can occur by multiple, sometimes overlapping, mechanisms including DNA binding proteins with lower specificity that work in combination with each other or other co-factors such as chromatin-associated long noncoding RNA(4). The complex nature of PRC2 recruitment in mammals, often necessitating multiple molecular mechanisms for action, has made it difficult to establish basic rules for PRC2 action, especially in *de novo* initiation of H3K27me3-triggered heterochromatin.

The finding that PRC2 binds with significant affinity to nearly every RNA tested in vitro(5,8,9), which then inhibits nucleosome methylation, has called into question the specificity of lncRNAs that were suggested to work with PRC2 to silence chromatin. However, these findings did help establish that the RNA being produced by an RNA polymerase on chromatin, nascent RNA, would be an intrinsic inhibitor of PRC2 activity(8), preventing initiation of heterochromatin at active genes. In fact, tethering a high-affinity RNA substrate for PRC2 directly to chromatin in the nucleus actively antagonizes PRC2 activity at normal target genes(10), due to the higher affinity of PRC2 for single RNAs with G-tracts than for chromatin(7,11). It has been proposed that the binding of PRC2 to nascent RNA may allow the complex to sample the landscape of the genome, searching for a context where PRC2 activity is promoted(5,6,12). This may occur through encounter of H3K27me3, guiding PRC2 to pre-established heterochromatin and allowing PRC2 activity(13). For *de novo* heterochromatin formation, it is less clear how the nascent RNA-inhibited PRC2 is activated. One model suggests that transcription shut down is required for PRC2 activity(14). This model is supported by the demonstrated antagonism of PRC2 by histone modifications of highly-active genes and nascent RNA(8,15). While both Xist and HOTAIR can turn off transcription upstream of PRC2 activity(16,17), the association of both of these lncRNAs with chromatin promotes H3K27me3 at lncRNA target loci. Therefore, even if the gene is repressed and little nascent RNA is present, the lncRNA is at the chromatin locus. Since Xist and HOTAIR RNAs inhibit PRC2, turning transcription off does not resolve the issue that an inhibitory chromatin-associated RNA is present at the site of PRC2 activity.

RNA and chromatin compete for PRC2 binding(7,11). Specifically, the linker DNA between nucleosomes competes with RNA(7). At equimolar nucleotide concentrations, single-stranded RNA wins the competition for PRC2 binding over chromatin(7), which explains the inhibitory effect of RNA for PRC2 activity. The affinity of PRC2 for a single-stranded RNA is much higher than for a hairpin version of the same nucleotide content(18), suggesting a more favorable context for chromatin to compete. Within the PRC2 mechanism of sampling chromatin via nascent RNAs and associating with lncRNAs on chromatin, we hypothesize there may be a context for double-stranded RNA that may tip the balance from RNA to chromatin for productive PRC2 interaction.

Double-stranded RNA (dsRNA) occurs in the nucleus, resulting from multiple mechanisms including intra-and inter-molecular RNA-RNA interactions. LncRNAs can make intermolecular RNA-RNA interactions with different types of transcripts. For example, Xist pairs with its antisense RNA, TsiX, during X inactivation(19). We have previously shown that HOTAIR can interact directly with target gene RNA, such as JAM2, through an imperfect RNA-RNA base-pairing match sequence(20). This specific matchmaking region within the sequence of HOTAIR is predicted to have a propensity for stable RNA-RNA interactions with known target transcripts(21). In fact, the region is a “hotspot” for RNA-RNA interactions as identified by computational prediction querying HOTAIR against the entire mRNA transcriptome(22). Interestingly, this “hotspot” overlaps a region within HOTAIR that was found to be conserved across vertebrates (except teleosts) with potential for RNA-RNA interaction with a HOXD transcript(23). HOTAIR RNA-RNA interaction was discovered due to its association with the RNA binding protein (RBP), heterogenous nuclear ribonucleoprotein (hnRNP) B1. Importantly, hnRNP A2/B1 was found to regulate HOTAIR-dependent PRC2 activity in cells. Furthermore, the B1 isoform was found to bind preferentially to HOTAIR and its target transcripts over A2(20). This protein has two tandem RNA recognition motif (RRM) domains that can associate in a head-to-tail dimer, binding two RNAs in an anti-parallel nature(24), suggesting a mechanism to promote RNA-RNA base-pairing interactions. We have previously shown that B1 can promote RNA-RNA interactions between HOTAIR and an RNA from a target gene, JAM2, which suggested that RNA matchmaking has a role in chromatin regulation by PRC2(20) (Fig. 1A), though the underlying mechanism behind this has remained unclear. LncRNAs such as Xist and HOTAIR can adopt favorable structured states(25,26), presenting an additional challenge for any proposed intermolecular RNA matchmaking where intramolecular interactions occur.

**FIGURE 1.**
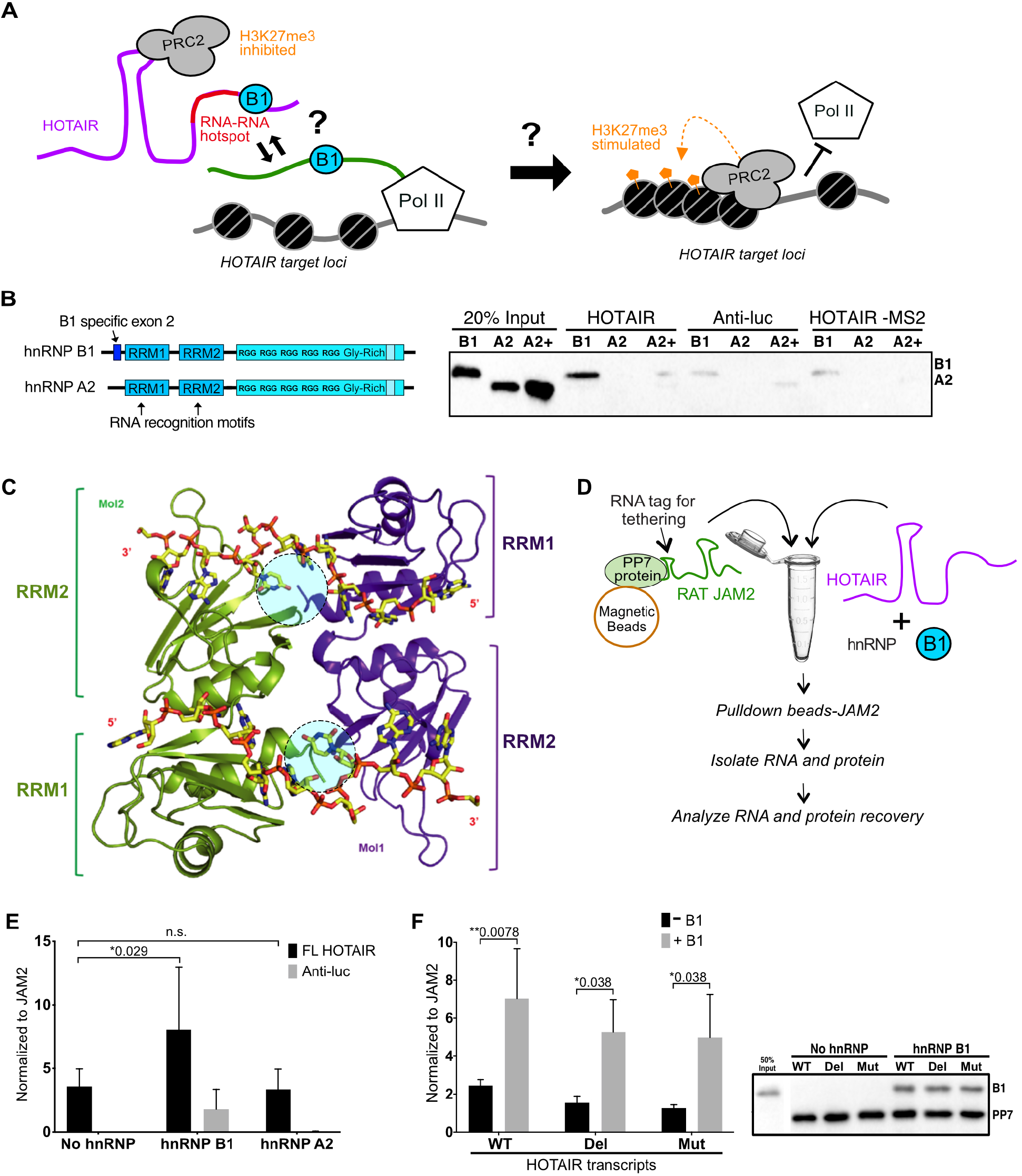
HOTAIR matchmaking is promoted by hnRNP B1. **(A)** Model of B1-mediated HOTAIR matchmaking to nascent target RNA leading to PRC2 activity and gene silencing, based on previous work. (**B)** Diagram of hnRNP B1 vs A2 domains, where B1 differs from A2 by inclusion of twelve unique amino acids encoding exon 2 (dark blue). In vitro RNA pulldown assays using in vitro transcribed MS2-tagged HOTAIR or Anti-luc with recombinant hnRNP B1 or A2. A2+ samples have 3X the amount of hnRNP A2 protein. Control samples with HOTAIR minus MS2-MBP fusion protein to account for background bead binding. Protein interaction with RNA was analyzed by Western blot with A2/B1 antibody. **(C)** Crystal structure of the tandem RRM domains of hnRNP A2/B1 in complex with 10 mer RNA(24). The RNA backbone is colored in yellow. Two molecules of the tandem RRMs are shown in purple (Mol1) and green (Mol2). The N-terminus, where the B1 NTD would protrude is highlighted with a blue circle. Figure adapted from (24). **(D)** Schematic representation of the in vitro RNA-RNA interaction assay (see Methods): IVT RAT-tagged JAM2 RNA and IVT HOTAIR is incubated in the presence or absence of recombinant hnRNP B1. RAT-tagged JAM2 was tethered to a PP7 coat protein which can then be isolated with magnetic beads. Recovery of HOTAIR by JAM2 pulldown was quantified by RT-qPCR and protein recovery by Western blot. **(E)** In vitro RNA-RNA interaction assays with RAT-tagged JAM2 RNA and either full-length HOTAIR or Anti-luc RNA in the presence or absence of recombinant hnRNP B1 or A2. The recovery of HOTAIR and Anti-luc RNA by JAM2 pulldown was quantified by RT-qPCR, normalized to JAM2 recovery (n=6). **(F)** In vitro RNA-RNA interaction assays with RAT-tagged JAM2 and either full-length HOTAIR, HOTAIR with the JAM2 interaction site deleted or the JAM2 interaction site mutated (to its complement base) in the absence or presence of hnRNP B1. The recovery of HOTAIR RNA with JAM2 was quantified by RT-qPCR and normalized to JAM2 recovery (n = 3). Protein recovery was analyzed by Western blot. (**E**,**F**) Error bars represent standard deviations based on independent experiments. Bar plots represent means. *P* values were determined using two-way ANOVA tests.

In the current study, we gain mechanistic insight into how B1-mediated RNA matchmaking can modulate HOTAIR structure and function to promote PRC2 activity. We determine the necessary features of hnRNP B1 for HOTAIR interaction and identify the specific regions of HOTAIR that interact with B1. We use chemical probing of HOTAIR secondary structure to highlight the structural changes that occur when B1 and an RNA-RNA match engage with HOTAIR. We find that genome-wide HOTAIR-dependent PRC2 activity occurs at loci whose transcripts make more-favorable RNA-RNA interactions with HOTAIR. Finally, we demonstrate that specific intermolecular RNA-RNA interaction relieves the inhibitory nature of HOTAIR RNA for PRC2 methyltransferase activity on nucleosomes. By dissecting molecular details of RNA matchmaking within HOTAIR, we highlight a switch that can result in PRC2 recruitment and activation by a lncRNA on chromatin. This may be a general mechanism that applies to many contexts where RNA plays a role in potentiating PRC2 activity.

## RESULTS

### RNA-RNA interaction is directly promoted by hnRNP B1 but not the A2 isoform

We have previously identified hnRNP A2/B1 as an important component of HOTAIR-dependent PRC2 activity in breast cancer. We found B1 was enriched preferentially in RNA pulldown assays with HOTAIR using nuclear extracts and we subsequently demonstrated direct *in vivo* binding of hnRNP B1 to HOTAIR, over the highly abundant isoform hnRNP A2. Additionally, B1 also bound preferentially to HOTAIR target transcripts, over A2(20,21). B1 differs from A2 by the inclusion of exon 2, which encodes 12 unique amino acids on the N-terminus (Fig. 1B). To further profile the recognition mechanism of HOTAIR by hnRNP B1 we first tested whether the isoform preference, B1 over A2, that was observed in the nuclear extract pulldown was recapitulated with purified protein. Using recombinant A2 or B1 proteins, expressed in *E. coli*, we performed HOTAIR pulldown assays and found that B1 binds preferentially to HOTAIR compared to A2 (Fig. 1B). Even a three-fold increase in A2 concentration did not recapitulate the same level of binding as B1 to HOTAIR. Little binding was observed for the control non-coding RNA, of similar size and GC content, which corresponds to the anti-sense sequence of the luciferase mRNA (Anti-luc), which we have used previously as a control(20). We conclude from this that the B1-specific N-terminal domain (NTD), directly confers preferential binding to HOTAIR (Fig. 1B). The presumed position of the NTD, based on the N-terminus position in the A2 isoform crystal structure(24) would place the NTD in proximity to bound RNA (Fig. 1C). This suggests the B1 NTD itself directly interacts with RNA as an extension of the RRM, to increase specificity, affinity, or both. Next we evaluated B1 vs. A2 in the RNA-RNA interaction assay we have previously used to characterize matchmaking using the HOTAIR target mRNA JAM2. Briefly, the RAT-tagged JAM2 match RNA fragment was incubated with full-length HOTAIR or Anti-luc control RNA in the presence or absence of hnRNP A2 or B1, and then affinity purified via the RAT tag. The association of HOTAIR or Anti-luc with JAM2 was quantified by RT-qPCR (Fig. 1D). Consistent with previous results, B1 was able to stimulate significant HOTAIR interaction with the target gene JAM2 RNA over A2. Moreover, B1 was able to promote a minimal amount of RNA interaction between JAM2 and the Anti-luc control RNA, suggesting that B1 can form an intermolecular bridge between two RNA molecules indirectly, as seen in the crystal structure for A2 (Fig. 1E).

### B1 bridges RNAs in the absence of strong base pairing potential

The crystal structure of the A2 tandem RRMs bound to RNA revealed a head-to-tail dimer in complex with two RNAs(24). Each RNA crosses the RRM of one protomer to the next and the two RNAs are aligned in an antiparallel nature (Fig. 1C). Based off of this RNA arrangement, where the dimer binds two molecules through single-strand engagement by the RRMs, we asked whether the ability of B1 to promote interaction of HOTAIR with its targets is dependent on base-pairing between the RNAs. We used the HOTAIR RNA-RNA interaction assay described above to test this. Matching of HOTAIR with its target does not require B1 in vitro, as we have previously shown(20). In fact, addition of B1 in our previously published RNA matching assays only modestly promoted the interaction between the RNAs. To test whether B1 can promote interaction with RNAs that do not have strong complementarity, we reduced the concentration of the RNAs to promote more stringency and B1 dependence. Under these conditions, B1 stimulates HOTAIR interaction with the JAM2 target RNA roughly three-fold (Fig. 1F). When the fairly extensive imperfect base-pairing interaction between JAM2 and HOTAIR is disrupted by deleting or mutating (changing to the complement base) the 64 nt interaction region on HOTAIR, B1 is able to recover nearly the same level of RNA-RNA interaction as with wildtype HOTAIR (Fig. 1F). This result suggests that B1 can bridge two RNAs due to the strength of binding to those individual RNAs and the ability of the dimer to interact with each RNA at the same time.

### Single-nucleotide mapping of B1 UV crosslinks to HOTAIR identifies major direct interactions

We used the eCLIP (enhanced Cross-Linking and Immunoprecipitation) method with recombinant B1 and in vitro transcribed HOTAIR to profile the interactions between the two(21). Briefly, UV crosslinking was performed on pre-incubated B1 and HOTAIR, the sample was digested with a low amount of RNase to generate RNA fragments that could be reverse transcribed and made into a sequencing library. The eCLIP protocol involves size selection and downstream ligation of one sequencing adapter to the 3’ end of the cDNA (Fig. 2A). Because a base that remains cross-linked to an amino acid “scar” can prematurely terminate reverse transcription, we mapped the termination sites of the eCLIP sequencing to better refine the specific site of direct B1 interaction on HOTAIR. The eCLIP results of B1 binding to HOTAIR revealed multiple regions with high reverse transcription (RT) stops. We observed six main peaks of RT stops from five locations on HOTAIR in the size of ∼25-100 nucleotides (Fig. 2B). Four of these peaks fell within domain 1 of HOTAIR (nucleotides 1-530), with the highest peaks at nucleotide positions 160, 310, 461-463 and 515, suggesting HOTAIR domain 1 is important for B1-mediated function (Fig. 2B). We conclude from this that B1 binds prominently to multiple distinct locations within domain 1 of HOTAIR. Based on other lower-frequency RT stops that still produce a peak, we suspect additional secondary interactions are made, potentially facilitated by proximity of RNAs in tertiary structure near the primary interaction sites.

**FIGURE 2.**
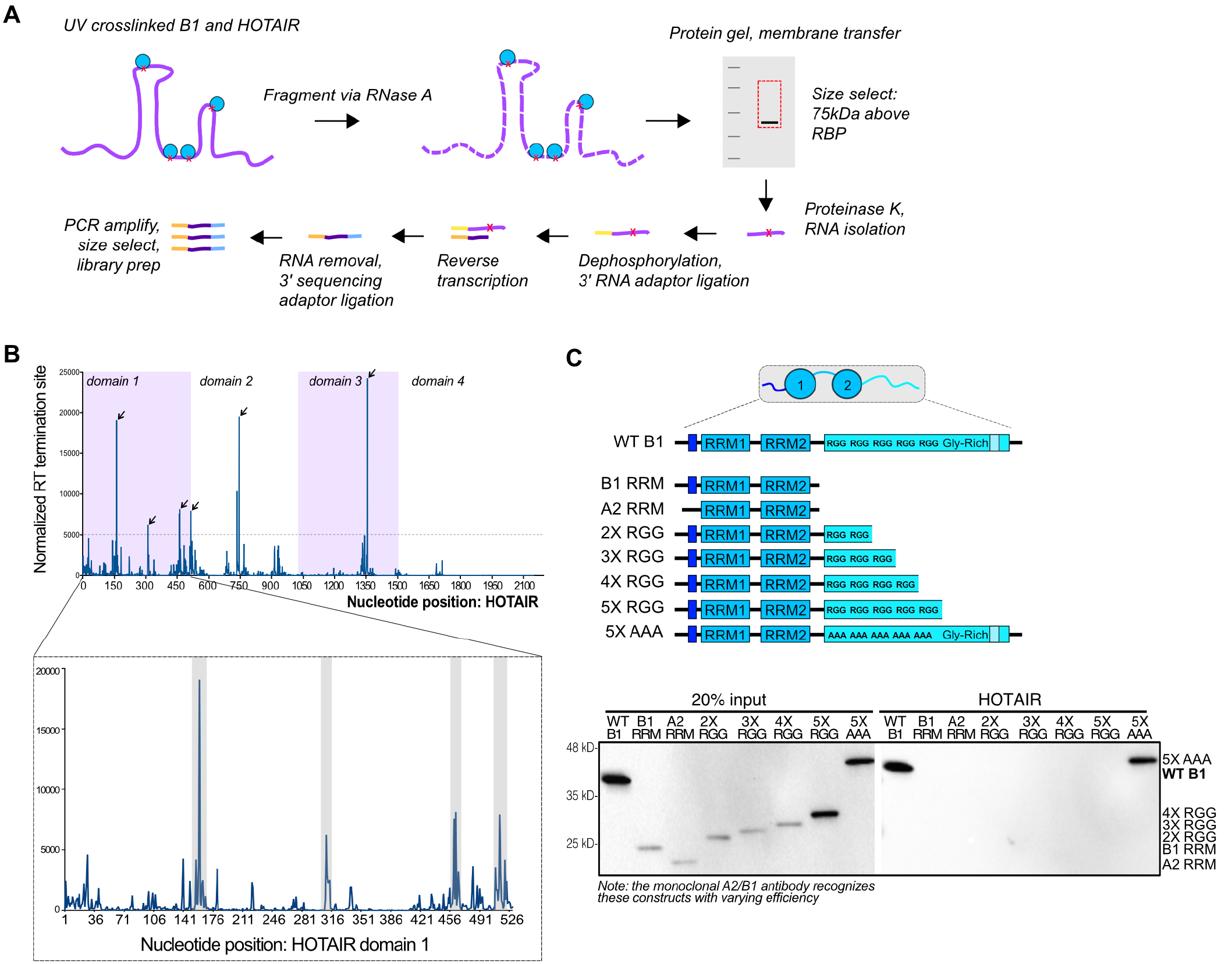
Examining hnRNP B1 specific interactions with the lncRNA HOTAIR using eCLIP mapping. **(A)** Schematic representation of the in vitro eCLIP experimental design: Recombinant B1 protein was incubated with IVT HOTAIR. Purified HOTAIR-B1 complexes were UV-crosslinked. RNA was fragmented with limited RNase A treatment, size-selected and processed for sequencing library preparation following the in vitro eCLIP protocol(21). **(B)** (*Top)* Mapping of reverse transcription termination events from HOTAIR-B1 in vitro eCLIP as a measure of direct protein crosslinking (HOTAIR domains alternately shaded violet). Termination sites were normalized to read count with significant peaks determined by values greater than 5000. (*Bottom*) Zoom in on domain 1 (1-530) highlights multiple B1 interaction sites (shaded in grey), based on termination site. **(C)** (*Top*) Diagram of hnRNP B1, B1 RRMs, A2 RRMs, B1 RGG truncations (2X-5X) and B1 RGG mutant (5X AAA). (*Bottom*) In vitro RNA pulldown assays using IVT MS2-tagged HOTAIR with recombinant truncated versions of hnRNP B1, A2 and a full-length B1 mutant with the five RGGs replaced with alanines (5X AAA). Protein association analyzed by Western blot for A2/B1. Intensities should be compared to input, since the monoclonal A2/B1 antibody recognizes the constructs differentially.

### B1 C-terminal glycine-rich domain participates in HOTAIR engagement

The crystal structure of the A2 RRMs demonstrates a minimal complex for bridging of RNAs (Fig. 1C). We have also demonstrated that the B1 N-terminal domain is required for efficient HOTAIR engagement and RNA matchmaking (Fig. 1B). Though these interactions are important features of RNA matchmaking, we wished to also test other features of B1 that may participate in this activity. In addition to the NTD and tandem RRMs, B1 has a run of “RGG” motifs proximal to the second RRM domain, as well as a low-complexity glycine-rich C-terminal domain. We generated minimal RRM constructs for B1 and A2, truncations eliminating RGGs and an alanine substitution of all five RGGs in the full-length construct. RNA pulldown experiments demonstrated a clear requirement for the C-terminal portion of B1, however the RGG motifs were unnecessary, suggesting the unstructured glycine-rich domain is required for tight binding to HOTAIR (Fig. 2E). The additional amino acids within glycine-rich domains have been proposed to influence the interplay of this region with other RBPs or RNA (27,28).

### Chemical probing highlights B1 interaction with HOTAIR

To further investigate mechanistically how RNA-RNA matchmaking interactions are facilitated, we performed chemical probing experiments on HOTAIR domain 1 with B1 and/or the JAM2 RNA match (62 nt) (Fig. 3A). RNA was chemically modified using 1-methyl-7-nitroisatoic anhydride (1M7) or DMSO, where 1M7 reacts with the 2’-hydroxyl of the RNA backbone to form adducts on accessible or flexible nucleotides, which primarily occur in single-stranded regions of the RNA (Fig. 3A). Adduct formation was quantified using primer extension via reverse transcription, where RT stops equate to reactivity to the modifier. Due to the strong RT stop introduced when HOTAIR and JAM2 interact, we had to include JAM2 removal steps to the protocol to generate complete reactivity data (Supplementary Fig. S3). The HiTRACE pipeline was used to analyze capillary electrophoresis data, and normalized reactivity values were evaluated for each condition: HOTAIR only, HOTAIR + B1, HOTAIR + JAM2, and HOTAIR + B1 + JAM2 (Fig. 3A,B). We observed reproducible chemical reactivity patterns across all conditions and were able to detect subtle and large changes in regions of HOTAIR upon the addition of B1 and/or JAM2 RNA consistent with our previous data and predictions for matchmaking (Fig. 3B,C). Interestingly, we found the addition of B1 reduced chemical probing reactivity in many regions of HOTAIR (Fig. 3B,C). This decreased reactivity was highlighted at the B1-interaction sites identified from our in vitro eCLIP analysis (highlighted in blue, 140-172 nt, 308-316 nt, 460-530 nt) (Fig. 3D,E). When we zoomed in on the specific B1 binding sites and compare them to the HOTAIR only condition, distinct attenuation of reactivity was observed within those regions, where the average change in B1 reactivity compared to HOTAIR alone (delta reactivity) was notably reduced (Fig. 3E). These results support a model of multiple prominent B1 interactions with HOTAIR and indicate that the eCLIP and chemical probing data are in agreement. Furthermore extent of reduced reactivity we observed may be explained by either 1) a broader direct influence of the protein on reactivity, perhaps through proximity in three-dimensional space, or 2) a significant change to RNA structure induced by B1, perhaps increased secondary structure, leading to decreased 1M7 reactivity in regions of HOTAIR that B1 does not bind directly.

**FIGURE 3.**
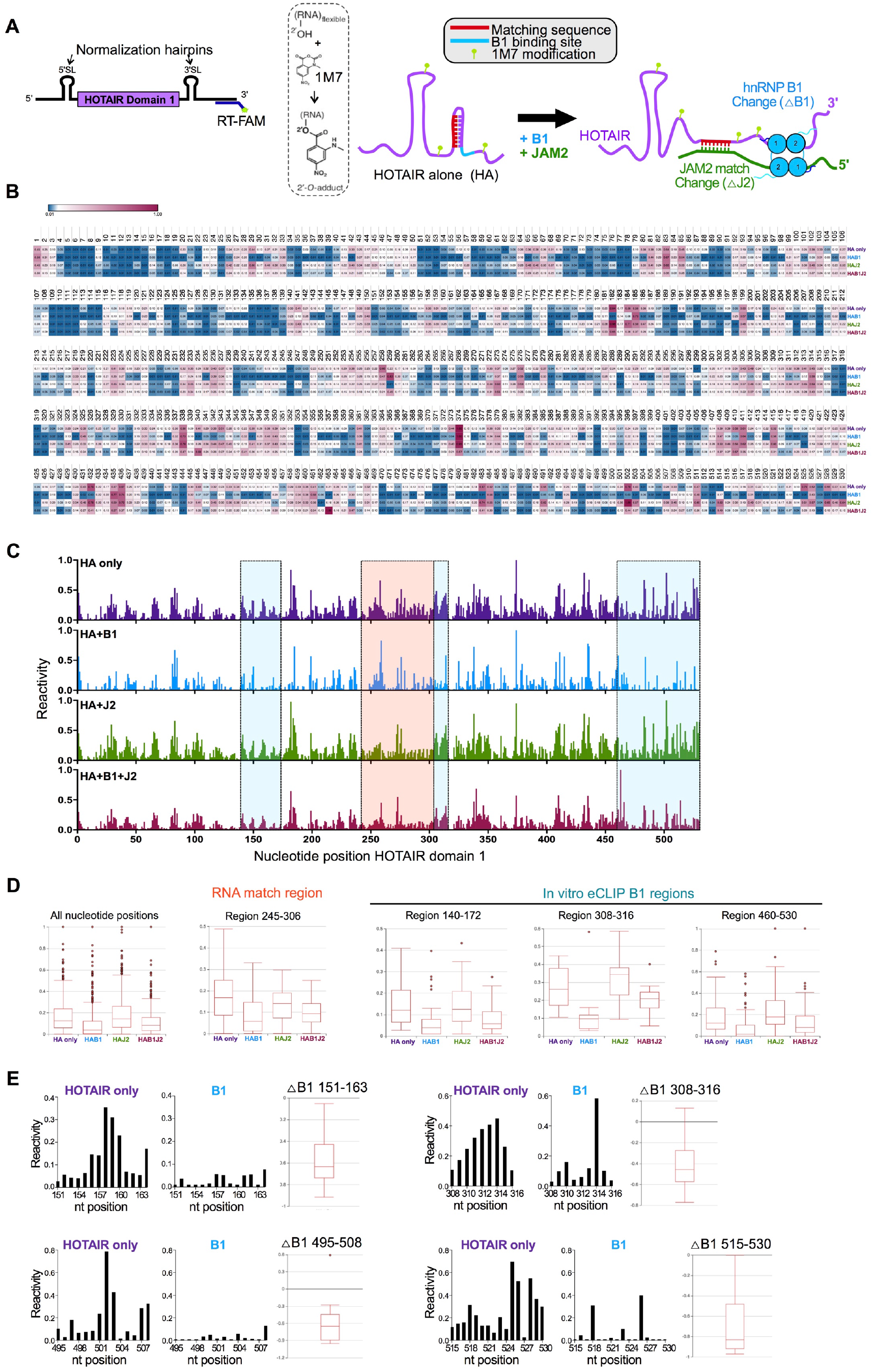
Chemical probing of HOTAIR highlights B1 interactions. **(A)** Diagram of the IVT HOTAIR domain 1 construct for chemical probing experiments. Schematic representation of 1M7 chemical probing of HOTAIR domain 1 with B1 and/or the JAM2 fragment (54 nt). **(B)** Heat map of normalized reactivity for HOTAIR only (HA only), HOTAIR + B1 (HAB1), HOTAIR + JAM2 (HAJ2), and HOTAIR + B1 + JAM2 (HAB1J2). Blue: less reactive, Red: more reactive. **(C)** Line graphs of normalized reactivity for each condition. B1 specific changes are highlighted in light blue and JAM2 changes in light red. **(D)** Boxplots of normalized reactivity for all nucleotide positions, the JAM2 interaction region (245-306 nt), and B1 eCLIP-derived binding sites (140-182 nt, 310-315 nt, 460-530 nt). **(E)** Bar graphs for normalized reactivity of HOTAIR only and HOTAIR + B1 at specific eCLIP B1 binding sites, by nucleotide. Boxplots to the right of each bar graph shows the cumulative change in reactivity compared to HOTAIR alone for HOTAIR + B1 (*△*B1) for the same region.

### RNA matchmaking alters HOTAIR structure

Next, we analyzed the change in HOTAIR 1M7 reactivities in the presence of the intermolecular RNA-RNA match with JAM2 (Fig. 4A). In general, we observed more regions with a gain in reactivity upon addition of JAM2 matching RNA. We defined specific regions of change by using the guidelines of five or more consecutive nucleotides with an average change in reactivity compared to HOTAIR alone (delta score) of 0.15 or higher and defined the boundaries by at least two consecutive nucleotides having 0.05 or less delta score (Fig. 4A). When we mapped these regions to the previously determined secondary structure model of HOTAIR domain 1(25), we found that the majority of the regions with higher reactivity upon JAM2 addition were opposing or adjacent to the region in HOTAIR that matches to JAM2 (Fig. 4B). These results suggest that binding of the JAM2 match alters HOTAIR conformation at these locations, potentially releasing them from intramolecular base-pairing, as would be predicted when JAM2 matches the other strand. In the presence of both B1 and JAM2, the reactivity profile in these regions is intermediate between B1 and JAM2 separately with HOTAIR, perhaps balancing the effects of protein or matching RNA alone, which may in turn promote RNA-RNA matchmaking activity. The JAM2 interaction site itself on HOTAIR (nucleotides 245-306) decreased in reactivity with JAM2 addition, with or without B1, likely because this region in HOTAIR maintains base-pairing, while transitioning from intramolecular to inter-molecular interactions (Fig. 4).

**FIGURE 4.**
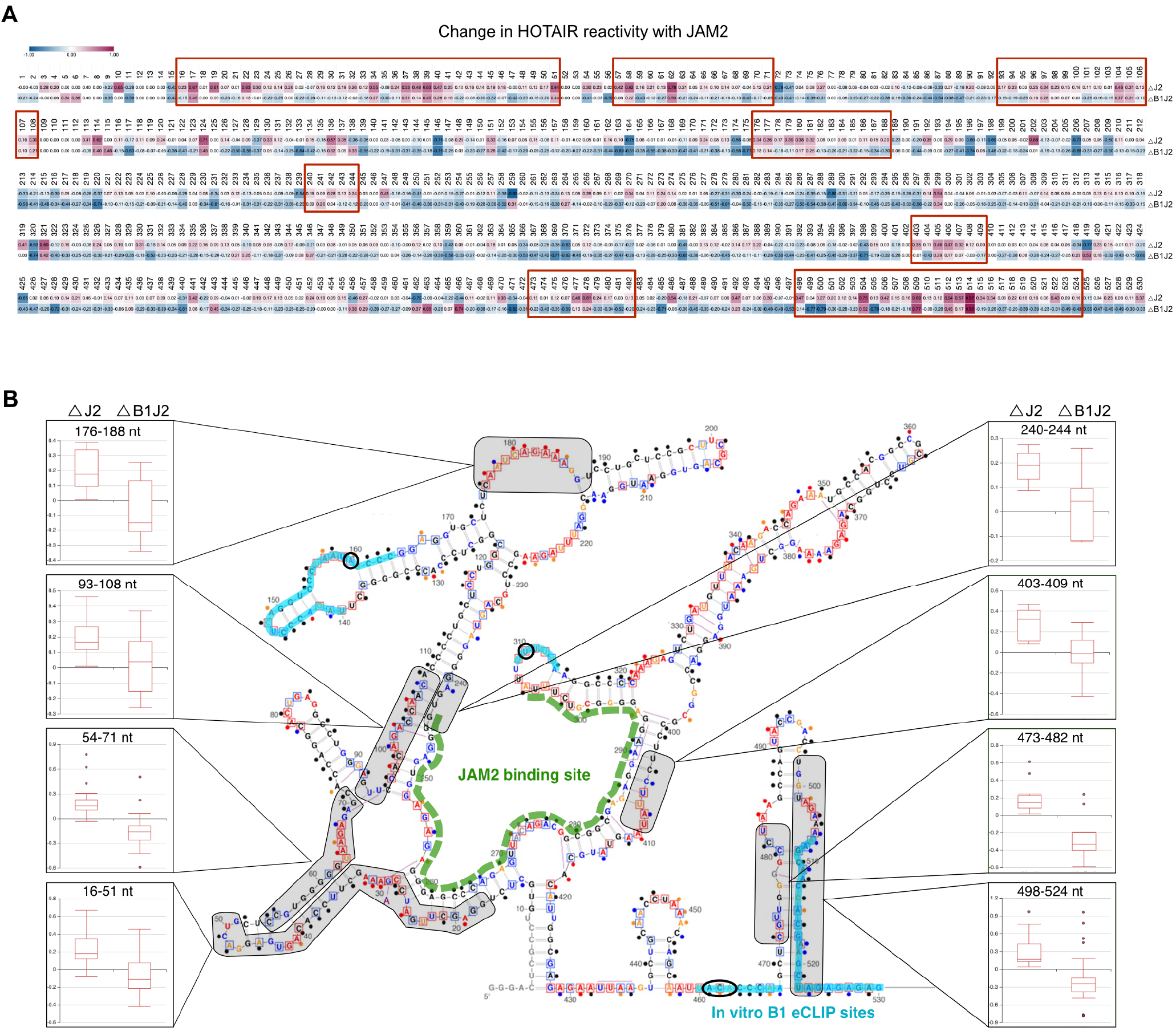
JAM2 matchmaking alters HOTAIR structure. **(A)** Heat map of HOTAIR reactivity changes upon addition of JAM2 (△J2) or both (△B1J2), compared to HOTAIR only. Key regions of higher reactivity produced by JAM2 interaction with HOTAIR are boxed in red. Specific regions of change were defined by using the guidelines of five or more consecutive nucleotides with a delta score of 0.15 or higher and defined the boundaries by at least two consecutive nucleotides with a delta score of 0.05 or less. Delta score is the average change in reactivity compared to HOTAIR alone, calculated using the formula [(HAJ2-HA)/(HAJ2+HA)]. **(B)** Boxplots of the altered regions from above, mapped to HOTAIR domain 1 secondary structure(25). In vitro eCLIP B1 binding sites are highlighted in blue and the JAM2 RNA-RNA interaction site is highlighted in green. Note that many of the regions of significant higher reactivity with JAM2 are base-paired or proximal to the JAM2-interacting region on HOTAIR (highlighted in grey).

### Conversion from ssRNA to dsRNA promotes PRC2 activity

It is not clear how PRC2 goes from RNA-mediated inhibition to an enzymatically active state in situations where a lncRNA like Xist or HOTAIR are known to be present and no pre-existing H3K27 methylation has occurred. PRC2 has been shown to have a strong affinity for ssRNA, but has a weaker affinity for duplex RNA, like that found in perfectly base-paired stem-loop structures(18). We analyzed RNA-RNA interaction predictions between HOTAIR and the entire transcriptome(21,22) and compared them to previously identified HOTAIR-dependent PRC2 targets in a model of HOTAIR-overexpressing triple negative breast cancer cells(20,29). Genes that acquired new PRC2 activity when HOTAIR was overexpressed were biased towards those with transcripts that have more-favorable RNA-RNA interaction propensity (Fig. 5A). This led us to hypothesize that duplex RNA might provide the correct context for PRC2 enzymatic activity by allowing PRC2 to transfer to chromatin via lower affinity binding of dsRNA. To determine this, we performed histone methyltransferase (HTMase) assays with single-stranded or double-stranded HOTAIR RNA and evaluated levels of H3K27me3. We first assessed optimal H3K27me3 activity in these assays using di-nucleosome templates composed of two 601 sequences surrounding 40-bp of linker DNA (Fig. 5B), recombinant PRC2 complex (Fig. 5C) and the co-factor JARID2 (Fig. 5D). We next annealed HOTAIR RNA with a titration of reverse complement RNA to form perfect duplex RNA and introduced these into the HTMase assays (Fig. 5E). We observed a distinct reduction in H3K27me3 in the presence of ssRNA vs no RNA. As we increased the amount of reverse complement RNA, H3K27me3 levels increased in a manner concurrent with formation of dsRNA (Fig. 5E). These results suggest that while a ssRNA inhibits PRC2 activity, formation of a duplex between the inhibitory RNA and its match is able to relieve this inhibition and promote PRC2-mediated H3K27me3.

**FIGURE 5.**
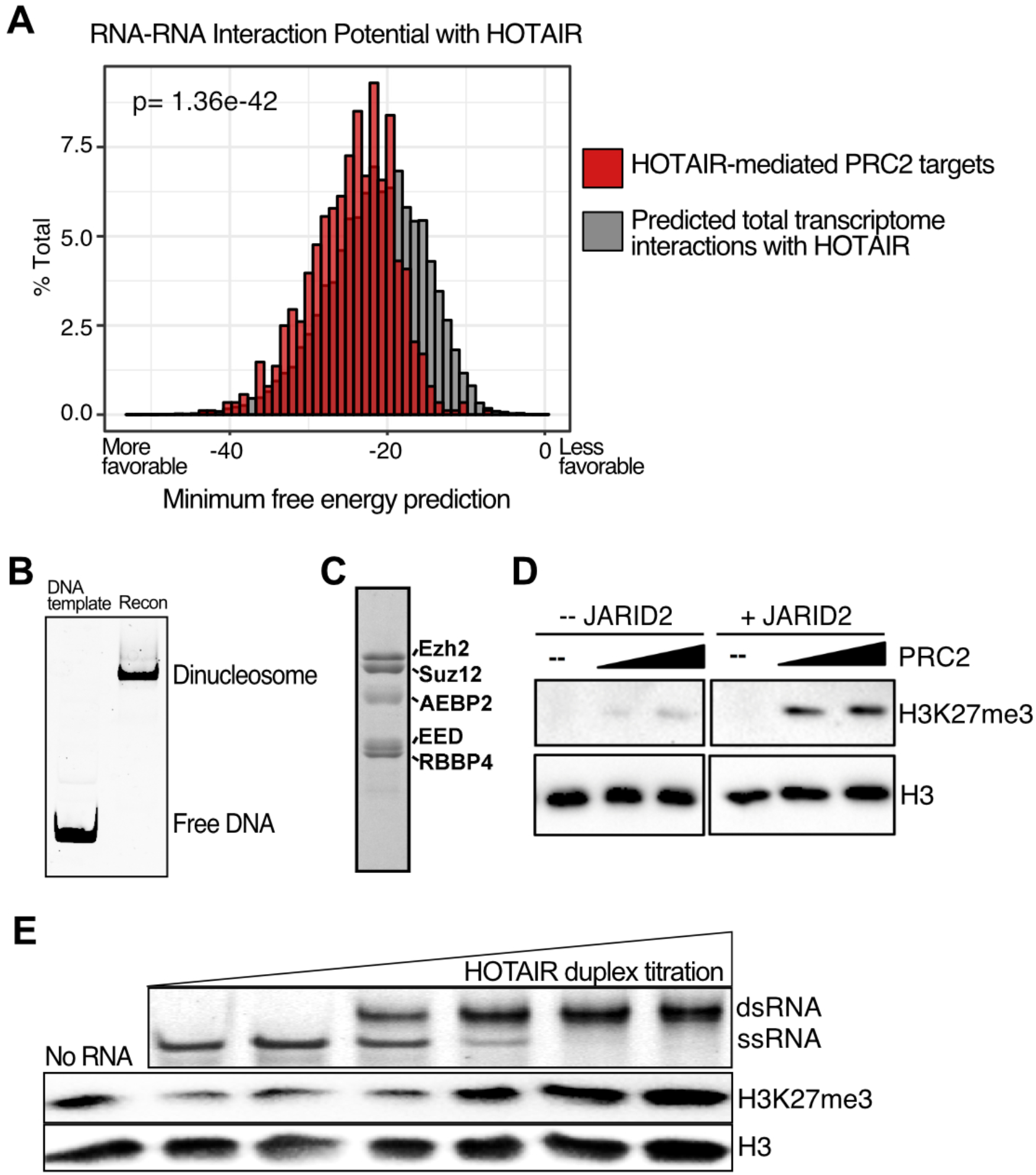
Duplex RNA promotes PRC2 activity. **(A)** Histograms comparing the predicted minimum free energy for RNA-RNA interactions between HOTAIR and 40,740 RNAs across the transcriptome (grey) or between HOTAIR and 885 transcripts from genes that gain PRC2 activity when HOTAIR is overexpressed in breast cancer cells (red). Data from(20,22,29). **(B)** Native gel of di-nucleosomes reconstituted via salt dialysis using a DNA template containing two 601 sequences surrounding 40-bp of linker DNA. DNA and nucleosome samples were run on a 5% native polyacrylamide gel and stained with SYBR Gold. **(C)** Recombinant human PRC2 complex includes SUZ12, EZH2, EED, RBBP4 and AEBP2, analyzed by SDS-PAGE and stained with Coomassie blue. **(D)** Histone methyltransferase assay (HMTase assay) was performed with recombinant PRC2 complex, di-nucleosomes, S-Adenosylmethionine (SAM) with and without the co-factor JARID2 (amino acids 119-574). PRC2 activity was determined by SDS-PAGE followed by H3K27me3 and total H3 Western blot analysis. **(E)** Native 0.5X TBE gel of RNA annealing titration with HOTAIR forward and reverse fragments to show formation of dsRNA. HMTase assay with annealed HOTAIR dsRNA titration.

### HOTAIR target RNA-RNA matches promote PRC2 activity

The experiments above used fully-duplex dsRNA, which may be relevant for some lncRNA contexts, but HOTAIR makes imperfect matches with RNA targets. To begin to address if HOTAIR imperfect RNA-RNA matches with target mRNA could also promote PRC2 catalytic activity, we evaluated the HOTAIR matches with endogenous targets JAM2 and HOXD10 (Fig. 6A). Consistent with previous results, HOTAIR alone was able to inhibit PRC2 activity. When increasing ratios of HOTAIR-JAM2 duplex were present, the match relieved the inhibitory effect of the single-stranded HOTAIR fragment alone, thereby stimulating H3K27me3 (Fig. 6B). Equal molar ratios of HOTAIR to JAM2 resulted in the highest levels of H3K27me3 promotion (Fig. 6C). Similar results were observed with the HOTAIR-HOXD10 duplex RNA match (Fig. 6D), with equal molar ratios of match RNA relieving PRC2 inhibition the most (Fig. 6E). In contrast, a control polyA RNA of similar size did not significantly relieve inhibition (Fig. 6F).

**FIGURE 6.**
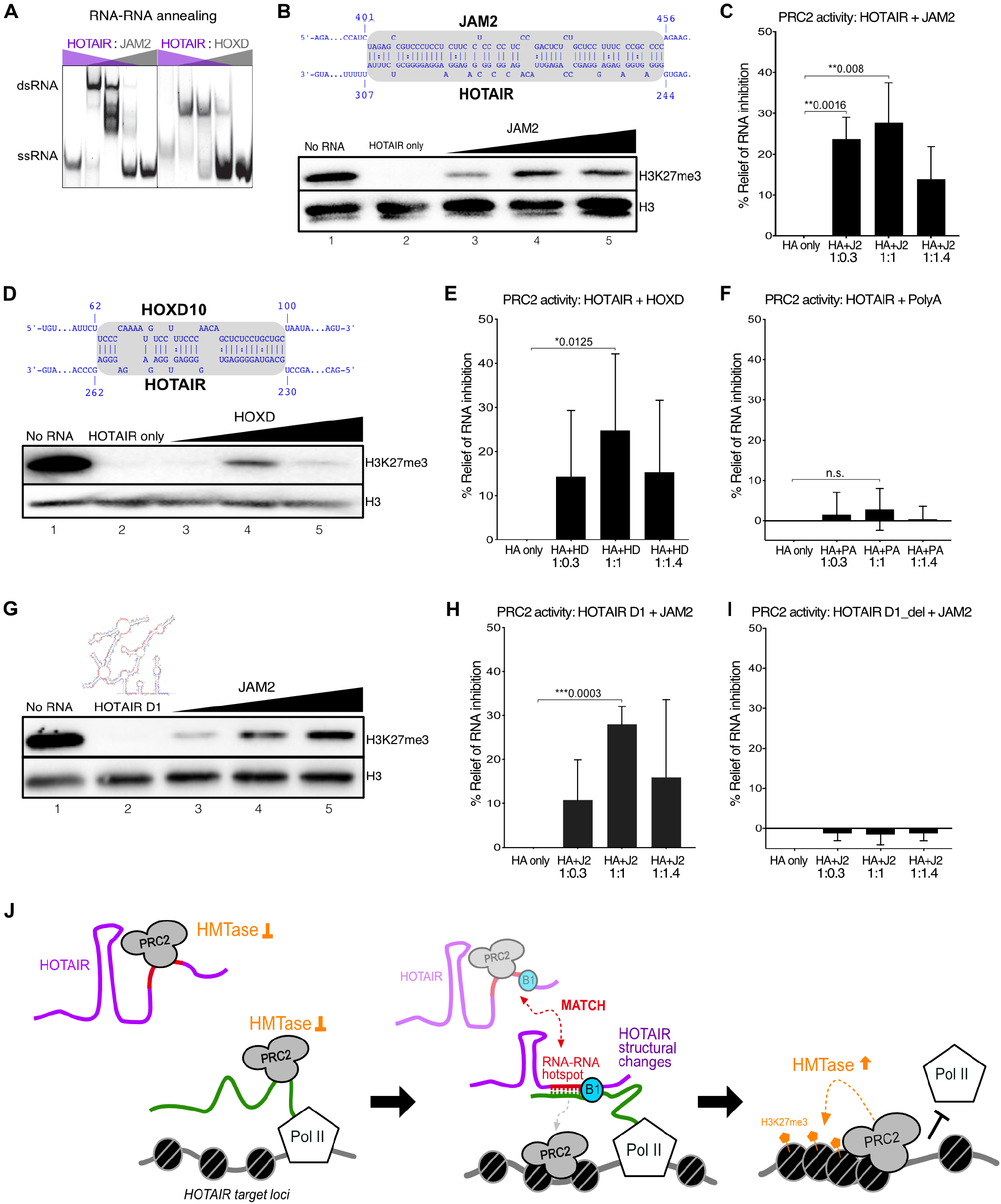
HOTAIR matching with target RNA promotes PRC2 activity. **(A)** Native polyacrylamide gel of RNA annealing titration with HOTAIR fragments and either JAM2 or HOXD10 matching fragments. **(B)** RNA-RNA interaction between HOTAIR and JAM2 predicted by IntaRNA(40). HMTase assay performed with recombinant PRC2 complex, di-nucleosomes, HOTAIR fragment (62 nt) and a titration of the JAM2 match RNA (54 nt) in ratios 1:0.3, 1:1, 1:1.4. PRC2 activity was determined by SDS-PAGE followed by H3K27me3 and total H3 Western blot analysis. **(C)** Quantification of **B** (n=3). **(D)** RNA-RNA interaction between HOTAIR and HOXD10 predicted by IntaRNA. HMTase assay performed as above with 10 µM HOTAIR fragment (31 nt) and a titration of the HOXD10 match RNA (37 nt) in ratios 1:0.3, 1:1, 1:1.4. **(E)** Quantification of **D** (n=5). **(F**) As in D, with polyA instead of HOXD10 (n=4). **(G)** HMTase assay performed as above with HOTAIR domain 1 (nucleotides 1-530) and a titration of a JAM2 match RNA (62 nt) in ratios 1:0.3, 1:1, 1:1.6. **(H)** Quantification of **G** (n=3). **(I)** As in G, except using HOTAIR with the JAM2 match site deleted “HOTAIR D1_del” (**C**,**E**,**F**,**H**,**I**) Signal for percent relief of inhibition is normalized to H3 signal and relative to no RNA and described as % relief from HOTAIR only reaction. Error bars represent standard deviations based on independent experiments. *P* values were determined using unpaired t tests with a 95% confidence interval. (**J**) Model for HOTAIR-mediated silencing: Binding of PRC2 to single-stranded regions of HOTAIR or nascent RNA inhibits enzymatic activity, B1 promotes RNA-RNA matching of HOTAIR with target nascent RNA via conformational changes in the lncRNA, thereby reducing PRC2 affinity for those RNAs through formation of the dsRNA match, initiating PRC2 interaction with chromatin leading to H3K27me3 and transcriptional repression.

The experiments above used RNA fragments surrounding the matching regions. These fragments were similar in size to the model RNA substrates used in previous biophysical studies of RNA-PRC2 interaction(18). Next we tested a much larger portion of HOTAIR RNA, using the HOTAIR domain 1 sequence (nt 1-530), to evaluate if the JAM2 match was sufficient to relieve inhibition of PRC2 activity. We found that HOTAIR domain 1 was a more-potent inhibitor of PRC2 by molar ratio, consistent with longer RNAs binding with higher affinity to PRC2. Strikingly, JAM2 was able to relieve PRC2 inhibition by this longer RNA (Fig. 6G,H). However, when we substitute HOTAIR domain 1 with a version in which the JAM2 interaction site is deleted, we observe no relief of inhibition (Fig. 6I). Together, these results demonstrate that the formation of dsRNA via HOTAIR matches with known genomic RNA targets like JAM2 and HOXD10, are sufficient to promote H3K27me3 in vitro. This suggests that endogenous RNA-RNA interactions between lncRNA and target nascent transcripts could facilitate PRC2 catalytic activity for gene silencing at specific genomic locations (Fig. 5A,6J).

## DISCUSSION

RNA binding to PRC2 inhibits its catalytic activity(8,9). It has remained unclear how this inhibition is relieved in contexts where RNA is present in a region of chromatin that has no previously-deposited H3K27 methylation, including lncRNA-associated loci and genes bearing nascent transcripts. Based on our previous observation that the HOTAIR lncRNA makes preferential interactions with hnRNP B1(20), a multi-valent RNA binding protein that promotes RNA-RNA interactions, we have further profiled the molecular basis of this mechanism and how it relates to the observation that HOTAIR can somehow promote PRC2 activity when overexpressed in cancers(29). We find that hnRNP B1 uses multiple domains to engage HOTAIR in a manner that can bridge it to a target gene RNA (Fig. 1-2). When HOTAIR matches with a target RNA, the lncRNA structure is remodeled to accommodate the base pairing (Fig. 3-4). We find that HOTAIR-mediated PRC2 targets make more favorable RNA-RNA interaction with HOTAIR (Fig. 5A). In turn, the formation of HOTAIR duplex RNA with its targets limits the ability of the lncRNA to inhibit PRC2 activity (Fig. 6). PRC2 binds to many individual RNAs in the nucleus(4), including nascent transcripts, presumably in an inactive state. Our results suggest a model where an RNA can be a positive effector of *de novo* PRC2 activity in a context where RNA-RNA interactions relieve PRC2 inhibition on chromatin (Fig. 6J).

### HOTAIR RNA matchmaking by hnRNP B1

The crystal structure of the tandem RRMs of hnRNP A2/B1(24) demonstrates a potential for two RNAs to be engaged by an A2/B1 dimer in an anti-parallel orientation. While the RRMs likely only bind single-stranded RNA, the adjacent RNA sequences are in a favorable orientation for base-pairing. This engagement is inter-molecular in the crystal structure and likely explains the ability of B1 to bring two RNAs together, even when the base-pairing potential between them is limited (Fig. 1F). There is potential for the same mode of engagement to work intramolecularly, as well, and this may underlie the multiple sites of direct interaction we observe for B1 on HOTAIR (Fig. 2B) and the ability of the protein to reduce chemical reactivity of multiple regions of HOTAIR (Fig. 3). The C-terminal domain of B1 is necessary for HOTAIR binding and thus promotion of RNA-RNA interactions (Fig. 2C). The C-terminus of the related hnRNP A1 can bind RNA(27,28) and includes an intrinsically-disordered domain that has been shown to self-associate and phase separate at high concentrations(30). Whether all of these properties are important for the mechanism we have characterized remains to be determined; however, the ability to self-associate, at least into a dimeric state, would potentially promote the RRMs of two monomers forming the head-to-tail conformation that can promote RNA matchmaking (Fig. 2E).

### RNA-RNA interactions promote PRC2 activity

The ability of an RNA with base-pairing potential to relieve the inhibition of PRC2 that is imposed by a single RNA binding event may apply beyond the HOTAIR mechanism. There are multiple examples of lncRNAs with intermolecular RNA-RNA interaction capability involved in PRC2 activity. For example, some imprinted loci such as *Kcnq1* have antisense transcripts that induce PRC2 activity and repression occurring coincident with sense transcripts, present in an RNA “cloud” at the locus that is methylated by PRC2(31). Xist also has a perfect complement antisense transcript, TsiX(31). In mice, Tsix transcription promotes PRC2 activity at the Xist promoter, coincident with the formation of Xist:TsiX double-stranded RNA, to repress Xist expression on the active X chromosome(19,32). In fact, this Xist:TsiX dsRNA does not inhibit PRC2(9). These RNA matching capabilities may underlie how PRC2 can methylate chromatin in a cloud of a lncRNA that would otherwise repress methyltransferase activity.

LncRNA matching may occur with the nascent transcript at the locus where PRC2 activity is promoted. Although nascent transcription from highly active genes is likely to “win out” by inhibiting PRC2(5,8), there are multiple pieces of evidence suggesting that PRC2 is active in the presence of nascent transcripts at lowly-expressed genes. Xist and Kcnq1ot1 lncRNAs are both present with low levels of their antisense transcript (TsiX and the protein-coding gene Kcnq1, respectively) while PRC2 methylates the chromatin at these loci(19,33). More-generally, H3K27me3 deposition has been shown to occur in genes with moderate transcription activity(34-36). Additionally, building up paused RNA Polymerase II with the drug DRB leads to more accumulation of promoter H3K27me3 than does complete inhibition of transcription initiation(14). Interestingly, a recent study found that endonucleolytic cleavage of nascent transcripts by a Polycomb-associated enzyme complex is important for maintaining low expression of Polycomb-repressed genes, suggesting nascent transcription persists even after PRC2 activity occurs(37). These results suggest that PRC2 can deal with the inhibitory effects of a nascent transcript to achieve H3K27 methylation. Intermolecular RNA-RNA interactions are one potential mechanism to achieve this. In addition, recent work has highlighted that once H3K27me3 has been established, this modification can help further relieve inhibition of methyltransferase activity by RNA(13). Our findings fit into a model where the inhibitory effects of RNA on PRC2 catalytic activity can be overcome by specific intermolecular RNA-RNA interactions to promote de novo H3K27 methylation. This mechanism may operate with multiple lncRNA pathways in normal contexts and when aberrantly high lncRNA expression occurs in disease, such as with HOTAIR, that may drive improper PRC2 activity.

## ACKNOWLEDGMENTS

We thank Kaushik Ragunathan, Srinivas Ramachandran, and members of the Johnson Lab for helpful comments on the manuscript. We thank Chen Davidovich and Tom Cech for PRC2 baculovirus constructs.

## Author Contributions

M.M.B and A.M.J conceived the study. M.M.B, E.W.H, C.B, S.K.W performed experiments. M.M.B, E.H.W, and J.T.R performed bioinformatic analysis. R.B. and A.M.G. designed and purified protein constructs and provided experimental advice. M.M.B, E.W.H, J.S.K, and A.M.J wrote the manuscript. All authors reviewed and approved of the manuscript.

### Data and Materials availability

All data related to the manuscript are included in main and supplemental materials. Specific materials generated during this study are available upon request.

## MATERIALS AND METHODS

### In vitro transcription of RNAs

In vitro transcription of RNA was performed using the MEGAScript T7 kit (Thermo Fisher Scientific or Ambion), reactions were incubated for 4 hours at 37°C. RNA was treated with Turbo DNase for 15 minutes, and RNA was purified with the RNeasy kit (QIAGEN).

### In vitro RNA pulldown experiments

15 nM of IVT 10X MS2 tagged RNA (Full-length HOTAIR or anti-Luc) was rotated (end-over-end) at room temperature for 15 minutes with 80 nM of recombinant hnRNP B1 or A2 in EMB 300 Buffer (10 mM Hepes pH 7.9, 300 mM NaCl, 3 mM MgCl2, 0.5% NP-40, 10% Glycerol, 0.1 mM PMSF, 0.5 DTT), RNase inhibitor (NEB) and 20 ug of competitor yeast tRNA (Roche) in a total volume of 300 uL per sample. At the same time, 300 nM MS2-MBP was prebound to 20 μL of amylose resin (NEB) in EMB 300 Buffer, RNase inhibitor (NEB) and 1% BSA, and rotated at room temperature for 15 minutes. The MS2-MBP amylose resin was then added to each IVT-hnRNP sample and incubated an additional 15 minutes rotating at room temperature. Resin was washed 4X in 800 uL EMB 300 Buffer and then protein association was analyzed by Western blot using antibody for hnRNP A2/B1 (Abcam #ab6102). Additionally, 10% of each sample was used for RNA analysis, where RNA was isolated by phenol/chloroform extraction, purified by ethanol precipitation, and quantified by RT-qPCR.

### Purification of PRC2 complex using Baculovirus expression system

Human PRC2 was purified essentially as described(5), using individual pFastBac1 constructs of EZH2, SUZ12, EED, RBBP4, and AEBP2 with HMBP-PrS (6XHis, MBP, and Prescission Protease sequences) N-terminal tags (courtesy of C. Davidovich and T. Cech). Briefly, equal MOI of each viral construct was used to infect Sf9 cells for 72 hours. at 27°C. Cells were harvested in PBS and snap frozen. Cells were thawed and resuspended in Lysis Buffer (10 mM Tris-HCl pH 7.5, 250 mM NaCl, 0.5% NP-40, 1 mM TCEP, 1X Complete Protease Inhibitor (Roche)), 20 mL per 500 mL culture, and slowly rocked for 30 minutes at 4°C. All further steps done at 4°C, but never on ice. Lysate was clarified at 29,000 RCF for 30 minutes. Supernatant was incubated with 0.75 mL amylose resin (NEB) for 2 hours. Sample was poured into a column and resin was washed with 8 mL Lysis Buffer, 12 mL Wash Buffer 1 (10 mM Tris-HCl pH 7.5, 500 mM NaCl, 1 mM TCEP), 12 mL Wash Buffer 2 (10 mM Tris-HCl pH 7.5, 150 mM NaCl, 1 mM TCEP). Protein was eluted with Wash Buffer 2 + 10 mM maltose. 0.5 mL fractions were collected until minimal protein was detected. Protein was concentrated ∼10-fold. Sample was incubated with PreScission Protease (GE Healthcare) overnight. Full PRC2 complex was separated by size-exclusion chromatography on a 24 mL Superose 6 column (GE Healthcare) in 10 mM Tris-HCl pH 7.5, 250 mM NaCl, 1 mM TCEP. Five-subunit complex containing fractions were concentrated, glycerol was added to 10%, and sample was aliquoted and flash-frozen.

### Recombinant protein purification in E. coli

Human hnRNP A2, B1 and B1 truncations were performed as previously described(20). Human GST-JARID2 (119-574) was expressed in BL21 (DE3) CodonPlus E. coli cells overnight at 18°C. Cells were lysed on ice using 2 mg/mL lysozyme and sonication in PBS (500 mM NaCl total), 1 mM EDTA, 1 mM EGTA, 1 mM PMSF, 0.5% Triton-X 100, 15 mM DTT. Sample was spun at 29,000 RCF at 4°C and supernatant was incubated with rocking with Pierce Glutathione Agarose Resin (Thermo Fisher) for 2-3 hours. Resin was washed with at least 20 times volume of PBS (350 mM NaCl total), 1 mM EDTA, 1 mM EGTA, 0.1 mM PMSF, 0.5% Triton-X 100, 0.1 mM DTT. Protein was eluted in 50 mM Tris-HCl pH 8.0, 350 mM NaCl, 1 mM DTT, 10 mM glutathione. Sample was dialyzed in 50 mM Tris-HCl pH 8.0, 350 mM NaCl, 1 mM DTT, 5% glycerol, aliquoted and flash frozen.

### In vitro RNA-RNA interaction assays

Performed as previously described (20) using in vitro-transcribed versions of HOTAIR, HOTAIR with JAM2 site deleted, HOTAIR with JAM2 site mutated (to its complement base) and the RAT-tagged JAM2 target as well as recombinant hnRNP B1 purified from E. coli.

### In vitro eCLIP-seq

In vitro eCLIP-seq was performed as previously described(21). 1.78 pmol HOTAIR RNA and 8.9 pmol recombinant hnRNP B1 (see (20) for sequence details) were incubated in a 1:5 RNA:protein molar ratio in 100 µL RNA refolding buffer (20 mM HEPES-KOH pH 7.9, 100 mM KCl, 3 mM MgCl_2_, 0.2 EDTA pH 8.0. 20% Glycerol, 0.5 mM PMSF, 0.5 DTT) for 20 minutes at room temperature. The mixture was diluted to 250 µL in refolding buffer and UV-crosslinked twice in one well of a 24-well plate at 250 mJ and 254 nm wavelength, with mixing by pipette in between. B1-RNA crosslinked samples were treated with 0.1 ng RNase A for 4 minutes at 37°C and 1200 rpm mixing, then stopped with 200 U Murine RNase Inhibitor (NEB). Following this, the *in vitro* samples were subjected to end repair, adaptor ligation, SDS-PAGE and transfer to nitrocellulose, and the remainder of the eCLIP-seq protocol, then sequenced multiplexed with other eCLIP-seq libraries as previously described(21). PCR products from different cycle number were analyzed by gel to avoid over-amplification (Supplementary Figure S1).

### Chemical probing and analysis

Chemical probing procedure was similar to(25,38). 20 pmol HOTAIR RNA was incubated in 500 uL reactions with equal molar JAM2 RNA, hnRNP B1 or both in RNA refolding buffer (50 mM HEPES pH 7.4, 200 mM KCl, 5 mM MgCl, 0.1 mM EDTA) at room temperature for 20 minutes. Reactions were divided into two tubes, each containing 245 ul. Reaction was started by addition of 544 nmol 1M7 (+) (1-methyl-7-nitroisatoic anhydride), or of an equal amount of pure DMSO as a control (–). Samples were incubated for 5 minutes at 37°C and reactions were quenched with 5 uL 0.5M MES-NaOH. Samples with the JAM2 RNA underwent a JAM2 removal step where they were incubated with JAM2 DNA complement oligo, heated at 94°C for 3 minutes to denature RNA and slow cool at room temperature for 10 minutes. All chemically modified RNA was purified using Trizol extraction + isopropanol precipitation and reverse transcribed using 0.2 uM RNA with SuperScript III reverse transcriptase (Thermo) at 48°C for 1 hour using a fluorescently labeled primer (IDT): 5’-/5-6FAM/. Labeled DNA products were eluted in HiDi formamide spiked Gene Scan ROX 1000 size standard (Thermo). Samples were run on an Applied Biosystems 3500 XL capillary electrophoresis system and the data was analyzed using HiTRACE RiboKit (38) with MatLab (Math Works). The HiTRACE normalized data are reported (supplemental table 2). HiTRACE normalized data for each condition was subsequently normalized to values between 0 and 1 to compare each condition on the same scale. In cases where 0.00 reactivity was called, the mean normalized value from each experimental condition was assigned a minimum possible value of 0.01 across all probing conditions (supplemental figure). This was done to avoid misleading high values when “fold changes” were calculated at positions of very low reactivity. Analysis of RNA chemical probing by regions was performed using Morpheus (https://software.broadinstitute.org/Morpheus) and comparative data where displayed back onto the previously determined RNA secondary structure model for HOTAIR (25).

### RNA-RNA interaction analysis of genome-wide PRC2 targets

Using a previously published database of computationally predicted interactions between human lncRNAs and the entire transcriptome(22), a list of 40,740 annotated mature transcripts and computational predictions for RNA-RNA interaction with HOTAIR were analyzed. This list was sorted by the predicted minimum free energy found among the interactions contained within each pair of RNA sequences. We compared the HOTAIR RNA-RNA interaction list with previously published HOTAIR-dependent PRC2 targets in MDA-MB-231 breast cancer cells identified by ChIP-seq(20,29), a combined list of 885 genes from multiple groups. Histograms displayed as a fraction of the total identified for each list were plotted together relative to predicted minimum free energy and a t test was performed comparing the two distributions.

### Nucleosome reconstitution

Di-nucleosomes were assembled using salt dialysis, as previously described(39). To generate a DNA template for chromatin reconstitution, a 343-bp PCR product consisting of 2x 601 positioning sequences separated by 40 bp of linker sequence was cloned into pUC57 vector backbone. PCR was purified using Macherey-Nagel Nucleospin DNA purification kit. Chromatin was reconstituted by salt dialysis: DNA template and human core histones were dialyzed 18-24 hours at 4°C from Hi salt buffer (10 mM Tris-HCl pH 7.6, 2M NaCl, 1 mM EDTA, 0.05% NP-40, 5 mM BME) to Lo salt buffer (10 mM Tris-HCl pH 7.6, 50 mM NaCl, 1 mM EDTA, 0.05% NP-40, 5 mM BME). Chromatin was dialyzed for an additional 1 hour in Lo salt buffer and concentrated using Ultra-4 10K Centrifugal Filter Device (Amicon) and stored at 4°C for no longer than a month.

### RNA annealing

RNA pre-anneal was performed by heating RNA at 94°C for 4 minutes in annealing buffer (6 mM HEPES pH 7.5, 60 mM KCl, 1 mM MgCl_2_), then slow cooling on bench for 40 minutes followed by placing samples on ice.

### Histone methyltransferase assays

HMTase assays were performed in a total volume of 15 μl containing HMTase buffer (10 mM HEPES, pH 7.5, 2.5 mM MgCl_2_, 0.25 mM EDTA, 4% Glycerol and 0.1 mM DTT) with 75 µM S-Adenosylmethionine (SAM, NEB), varying amounts ssRNA and duplex RNA (see above), 600 nM JARID2, 360 nM of dinucleosomes, and 600-660 nM recombinant human PRC2 complexes under the following conditions. The reaction mixture was incubated for 30 minutes at 30°C and stopped by adding 12 μl of Laemmli sample buffer (Biorad). After HMT reactions, samples were incubated for 5 minutes at 95°C and separated on SDS-PAGE gels. Gels were then subjected to wet transfer (30% MeOH transfer buffer) of histones to 0.22 μm PVDF membranes (Millipore) and protein was detected by Western blot analysis using primary *α*Rb H3K27me3 antibody (Millipore #07-449), secondary antibody (Biorad #170-6515), H3-HRP (Abcam #ab21054). Similar experiments were performed, except that the total ribonucleotide concentrations of all RNAs used were kept constant (Supplementary Fig. S2).

---

**SUPPLEMENTARY FIGURE 1.**
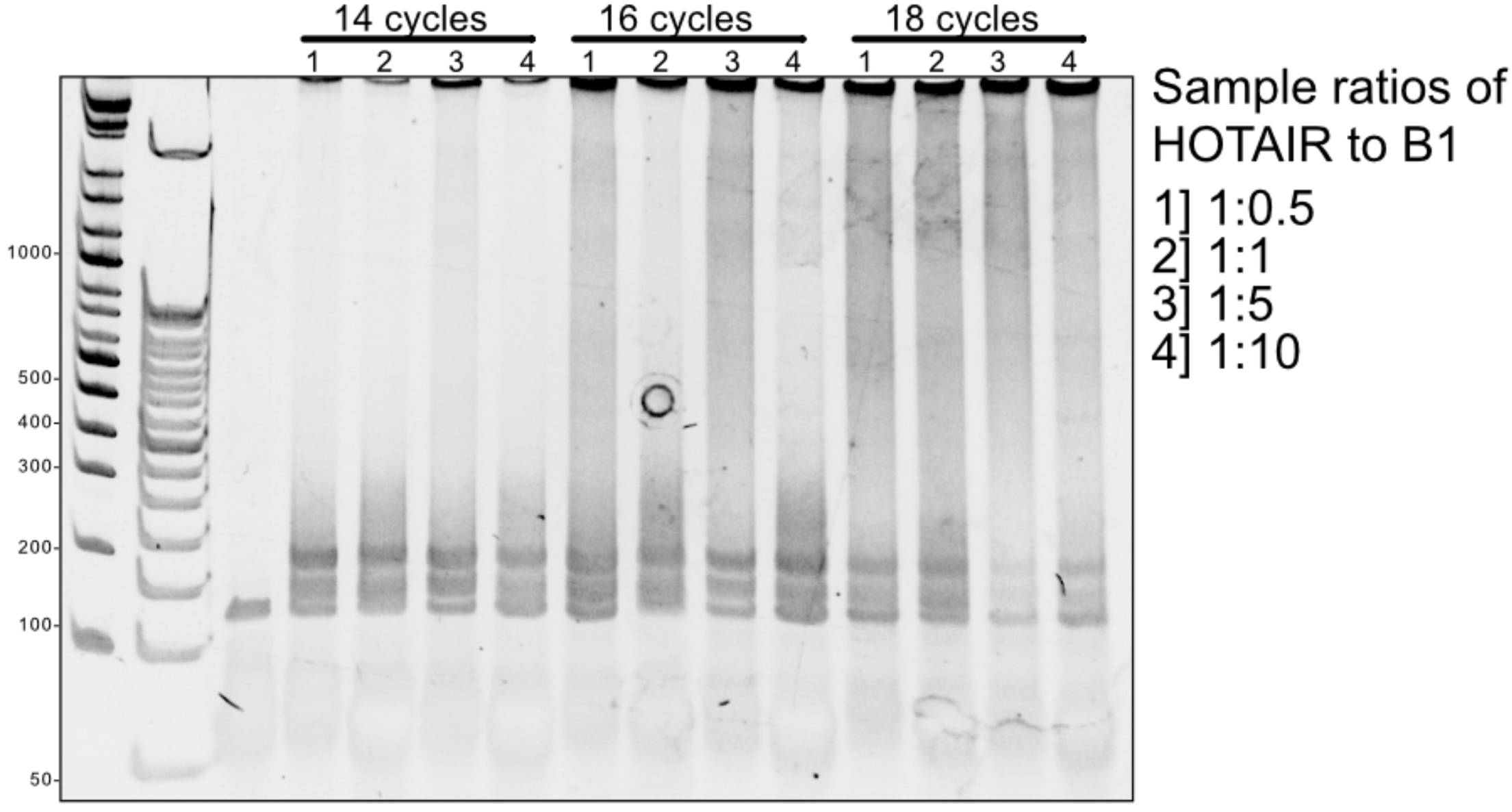
eCLIP PCR amplification of cDNA. Optimization of 1: 10 diluted cDNA amplification. PCR cycle number 14, 16 and 18 were tested based on the average qPCR Ct value for all samples. Cycle number 14 was selected based on product in the 100-300 range with minimal over amplification. Cycle number for final PCR is typically 3 cycles less then tCt of the 1: 10 diluted sample, so 11 cycles was used in the final PCR for cDNA amplification.

**SUPPLEMENTARY FIGURE 2.**
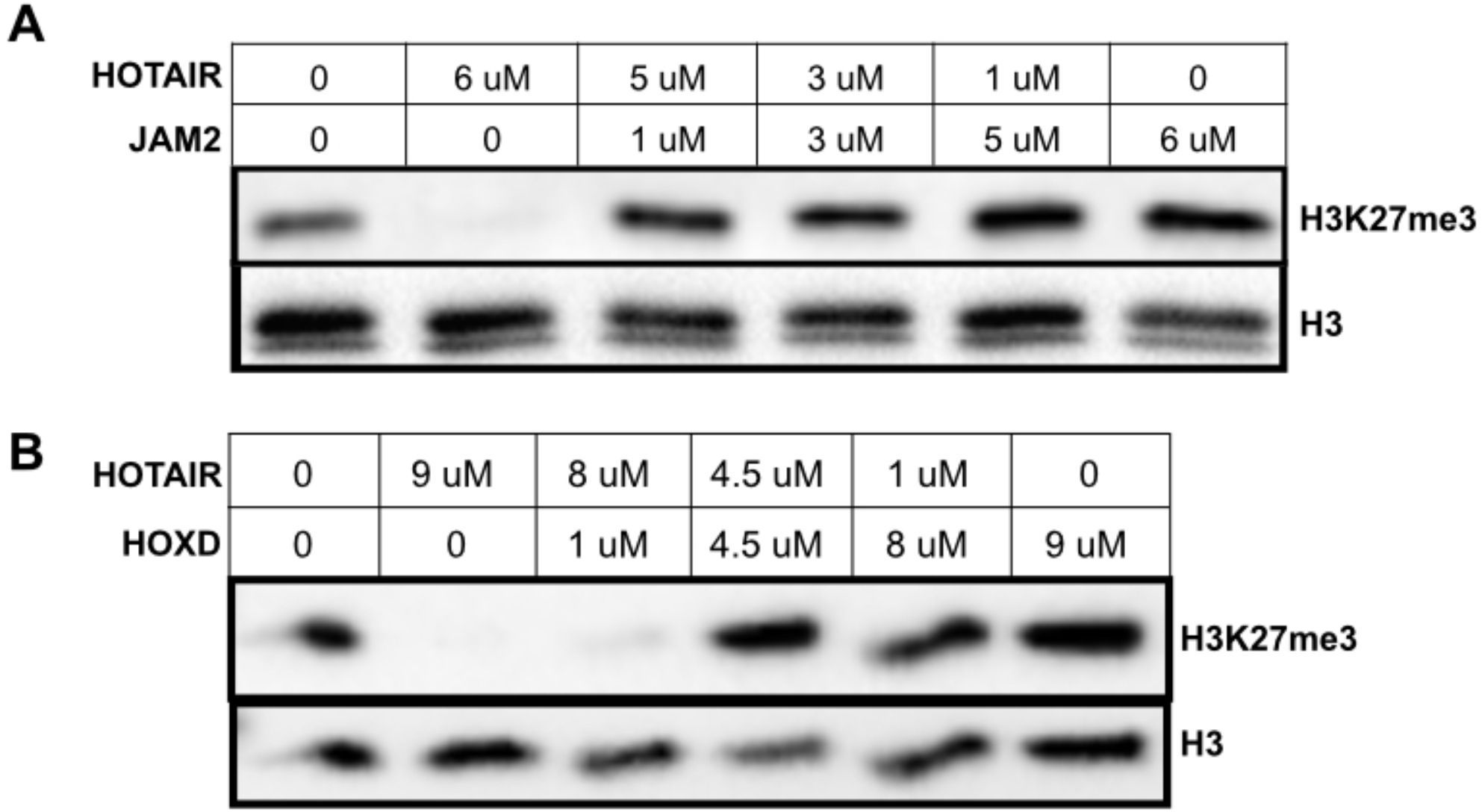
Control HTMase assays. **a,** HTMase assay performed with 650 nM of recombinant PRC2 complex, 360 nM di-nucleosomes, 75 uM SAM with titrations of RNA. Top blot had titrations of 6, 5, 3, 1, 0 uM HOTAIR-JAM2 match RNA with titration of 0, 1, 3, 5, 6 uM JAM2 match RNA. **b,**Bottom blot had titrations of 9, 8, 4.5, 1, 0 uM HOTAIR-HOXD match RNA with titration of 0, 1, 4.5, 8, 9 uM HOXD match RNA. H3K27me3 activity was determined by SOS-PAGE followed by Western blot analysis.

**SUPPLEMENTARY FIGURE 3.**
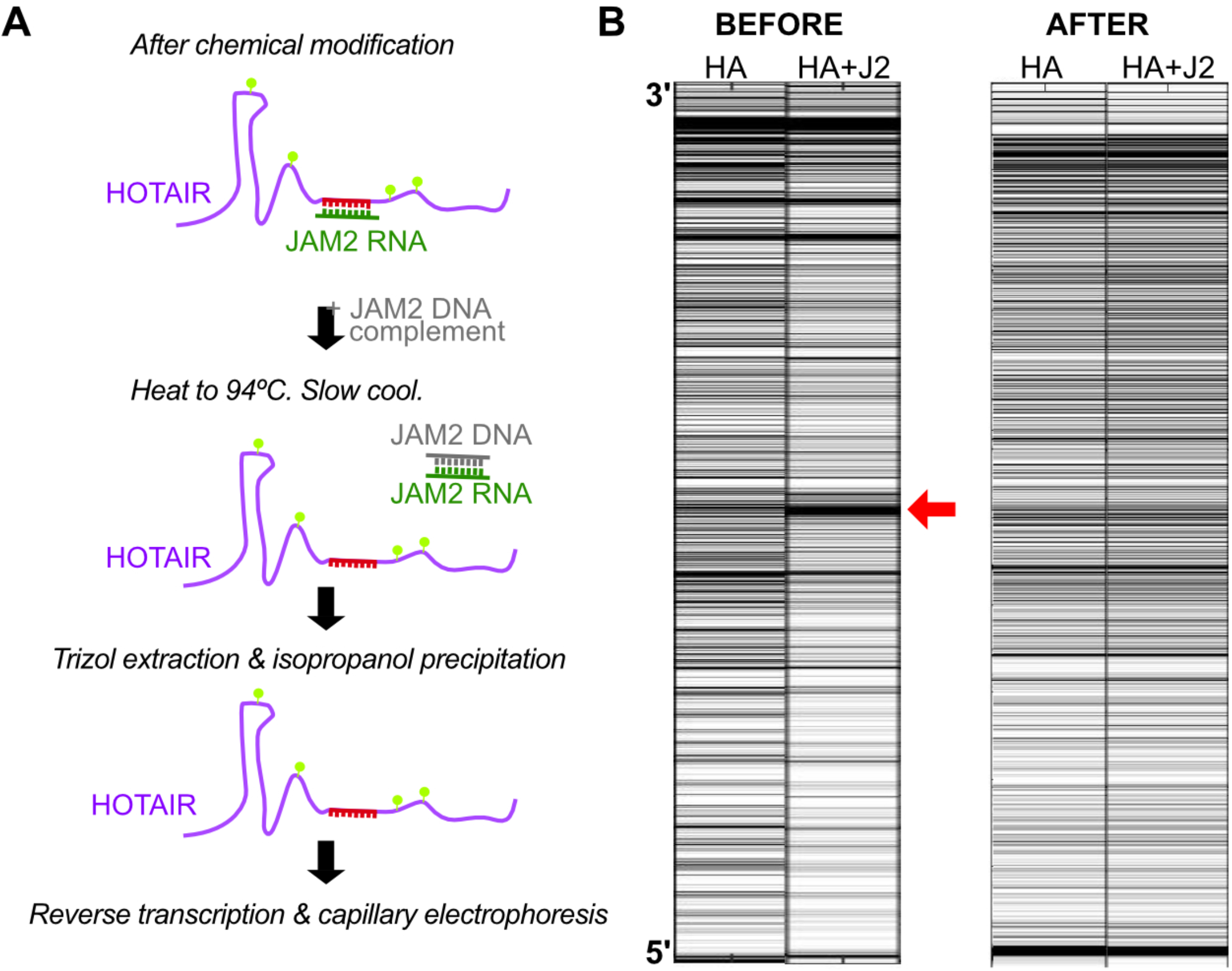
HiTRACE signal alignment before and after JAM2 removal steps added. **a,**Schematic of JAM2 removal steps. After chemical modification steps, a DNA complement of JAM2 is added and each sample is denatured at 94° C, then slow cooled. HOTAIR is then purified via Trizol extraction and isopropanol precipitation. Purified RNA is then reverse transcribed and analyzed by capillary electrophoresis. **b,**Signal channel data visualized as heatmaps for DMSO treated RNA samples with HOTAIR only vs HOTAIR + JAM2 before and after JAM2 removal steps were added to the protocol to reduce the strong RT stop induced by JAM2 RNA-RNA interactions with HOTAIR.

