## Supplemental Figures for "RNA matchmaking remodels lncRNA structure and promotes PRC2 activity"

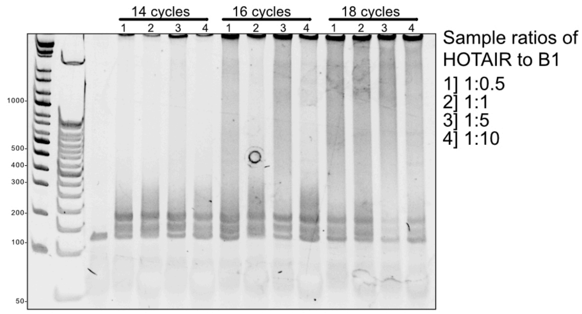


**SUPPLEMENTARY FIGURE 1. eCLIP PCR amplification of cDNA.** Optimization of 1:10 diluted cDNA amplification. PCR cycle number 14, 16 and 18 were tested based on the average qPCR Ct value for all samples. Cycle number 14 was selected based on product in the 100-300 range with minimal over amplification. Cycle number for final PCR is typically 3 cycles less then the Ct of the 1:10 diluted sample, so 11 cycles was used in the final PCR for cDNA amplification.

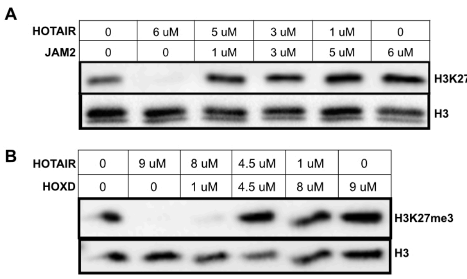


**SUPPLEMENTARY FIGURE 2. Control HTMase assays. a**, HTMase assay performed with 650 nM of recombinant PRC2 complex, 360 nM di-nucleosomes, 75 uM SAM with titrations of RNA. Top blot had titrations of 6, 5, 3, 1, 0 uM HOTAIR-JAM2 match RNA with titration of 0, 1, 3, 5, 6 uM JAM2 match RNA. **b**,Bottom blot had titrations of 9, 8, 4.5, 1, 0 uM HOTAIR-HOXD match RNA with titration of 0, 1, 4.5, 8, 9 uM HOXD match RNA. H3K27me3 activity was determined by SDS-PAGE followed by Western blot analysis.

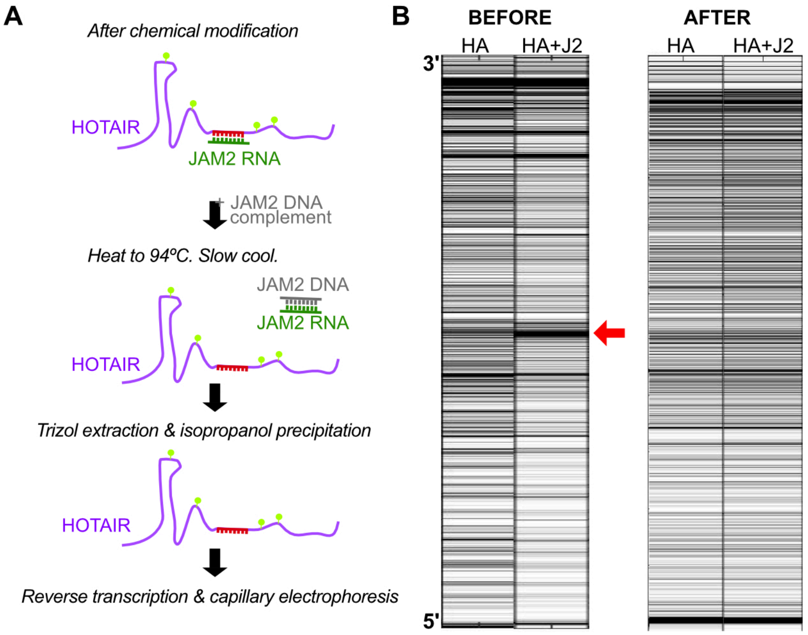


**SUPPLEMENTARY FIGURE 3. HiTRACE signal alignment before and after JAM2 removal steps added. a**,Schematic of JAM2 removal steps. After chemical modification steps, a DNA complement of JAM2 is added and each sample is denatured at 94ºC, then slow cooled. HOTAIR is then purified via Trizol extraction and isopropanol precipitation. Purified RNA is then reverse transcribed and analyzed by capillary electrophoresis.
**b**,Signal channel data visualized as heatmaps for DMSO treated RNA samples with HOTAIR only vs HOTAIR + JAM2 before and after JAM2 removal steps were added to the protocol to reduce the strong RT stop induced by JAM2 RNA-RNA interactions with HOTAIR.
